# How reliable are DAO supplements? — A comparison of over-the-counter Diamine oxidase products

**DOI:** 10.1101/2023.04.13.536689

**Authors:** Marc Alemany-Fornes, Jaume Bori, Maria Tintoré, Jordi Cuñé, Carlos de Lecea

**Author notes:** The authors contributed equally (Marc Alemany-Fornes, Jaume Bori).

## Abstract

Diamine oxidase (DAO) supplements have gained increasing attention in recent years due to their potential to support DAO deficiency, histamine intolerance and related symptoms. However, choosing a reliable and trustworthy DAO supplement can be a challenging task for patients and Healthcare practitioners. One of the main concerns is the lack of regulatory oversight on dietary supplements, which may result in misleading or incomplete labelling, incorrect dosages, and inadequate quality control. Such situations may lead to patients consuming supplements with insufficient amounts of active enzyme or with questionable purity, potentially resulting in undesired health outcomes. Thus, we tested the DAO activity of a variety of already in-the-market supplements and compared them with one another and against the specified activity, if any, by the manufacturer. Our results show a great discrepancy in most of the products and significant differences in DAO activity between different manufacturers.

## 1. Introduction

Diamine oxidase (DAO) is an enzyme that plays a crucial role in histamine metabolism, catalysing the oxidative deamination of histamine, putrescine, and other biogenic amines. DAO is primarily expressed in the small intestine and kidneys, and it contributes to the regulation of histamine levels in the human body(1).

Dietary supplement usage has experienced a significant increase over the last few decades(2–4). Evidence suggests that commercially available dietary supplements may not always contain the listed ingredients on the declared label. In addition, inconsistent quality control during manufacturing and preservation can lead to problems with the integrity and stability of the composition of supplements(5). Finally, some reports indicate that contamination levels can vary significantly not only between different production batches but also within a single package, indicating a lack of homogeneity in the substance content of these products. Consequently, some tablets or capsules may be uncontaminated, while others may test positive for contaminants(6,7).

Unfortunately, the current legislation does not make it mandatory for dietary supplements to be monitored for their composition with the same degree of severity as pharmaceutical products, hence raising concerns about their actual efficacy and safety. As patients increasingly rely on dietary supplements, it is crucial to assess their quality and credibility, especially taking into account that the aforementioned issues are exacerbated by the fact that most dietary supplements are readily available on alternative markets, such as online platforms. As a result, national supervisory bodies that exist to safeguard consumers may be circumvented by internet sales.

Recently, there has been an increasing interest and growing evidence in the use of DAO as a dietary supplement to support histamine metabolism, particularly among individuals with DAO deficiency and related disorders(8–11). In Europe, the commercialization of DAO-containing products is ruled by EU Regulation 2017/2470 which authorizes DAO to extract from pig kidneys as a novel food and accepts its use for the manufacturing of either Food Supplements (FS) or Food for Special Medical Purposes (FSMP). Within this regulation, the units applied to measure DAO activity are Histamine Degrading Units (HD)(12). However, it is noteworthy that manufacturers and researchers are increasingly adopting the International System of Units (SI) for measuring the activity of DAO supplements, which are usually expressed in U or mU per mg of extract (A unit of DAO activity is equivalent to the quantity of enzyme that can break down one micromole of substrate in one minute). This transition to the SI system permits facile comparison and harmonization of data in the scientific community as enzymatic reaction conditions are better defined than with HDU.

Despite the potential benefits of DAO supplementation, the quality and reliability of commercially available products remain a concern. Inaccurate labelling in product composition can compromise the effectiveness of DAO supplements and even pose potential health risks due to the consumption of rich histamine foods by patients under non-effective DAO supplementation. Therefore, it is critical to ensure that DAO supplements are produced and labelled accurately and that their efficacy and safety are supported by scientific evidence.

As a product of biological origin, it is a major challenge to ensure homogeneity in the manufacturing of DAO products. Previous studies have explored the variability of active ingredients within supplements containing ingredients from natural sources. Such variability can arise from the origin of the natural source (13) and the ontogenetic phase of the natural source at the time of retrieval (14). Furthermore, manufacturers may confront other intrinsic challenges, such as random effects, which difficult consistent products. Variations can inevitably occur during the manufacturing process, which underscores the necessity of implementing an appropriate quality control system and manufacturing process(15).

In this context, this study aimed to determine the diamine oxidase activity of DAO supplements by an enzymatic assay coupled to LC-MS/MS, using a reliable and already validated methodology, to provide further insights into the quality and consistency of commercially available DAO products.

## 2. Materials and methods

### a. Quantification of DAO activity

The determination of diamine oxidase (DAO) activity was performed by an enzymatic assay coupled to liquid chromatography-tandem mass spectrometry (LC-MS/MS), a methodology based on the one developed by Comas-Basté et al (16). To determine DAO activity in each sample, the absolute value of the slope of histamine consumption (30-120 min) measured in nmol was utilized to represent the activity in mU (nmol/min). Additionally, the weight of the porcine/vegetable extract was considered to determine DAO activity in mU/mg for each sample. The analysis was performed by Eurecat technological centre.

### b. Chemicals

Reference standards Histamine and Histamine-α,α,β,β-d4 were purchased from Merck and Toronto Research Chemicals, respectively. Diamine Oxidase from porcine kidney with lot number 011M7015 / SLCG5608 was also obtained from Merck. The enzyme was stored at -20°C until use. Prior to use, stock solutions were thawed and diluted in ultrapure water to prepare working standard solutions at the appropriate concentration.

Water, methanol and Acetonitrile LC-MS grade were obtained from Merck. Formic acid Ammonium formate and Perchloric acid were also purchased from Merck. Sodium dihydrogen phosphate and di-Sodium hydrogen phosphate were used to prepare the buffer for the enzymatic assay.

### c. Dietary supplements

Ten dietary supplements labelled to contain Diamine oxidase, have been used for this study. Seven of them listed porcine kidney extract as the source of Diamine oxidase and three of them are vegan versions, from which one is based on pea sprout extract, one contains peas and lentils extracts and the last one doesn’t specify the source of their DAO enzyme and does not provide any ingredient list, it just indicates “no major food allergens”.Detailed information can be found in Table 1.

**Table 1.** DAO supplements available in the market and their specifications. *The values for the amount of protein extract rich in DAO for the products Histamine manager and Histazyme have been estimated as the label of those products (600mcg=0,6mg) refers to the content of DAO enzyme, not to the content of protein extract, like the rest of products. For ease of comparison, 0.6 mg is the estimated content of DAO in 8.4 mg (2 capsules) of protein extract as stated in EU Regulation 2017/2470(17).

| Supplement name | Manufacturer | Active ingredient | Format | Intake recommended by manufacturer | Amount of active ingredient | Number of tablets per dose | Advertised Dao activity (HDU per dose) |
| --- | --- | --- | --- | --- | --- | --- | --- |
| DAOfood Plus | DR Healthcare | Porcine kidney extract | Capsules | 1 tablet: 3x day; before meals | 4.2 | 1 | 20,000 |
| DAOfood | DR Healthcare | Porcine kidney extract | Mini tablets | 1 tablet: 3x day; before meals | 4.2 | 1 | 20,000 |
| DAOfood Veg | DR Healthcare | Pea sprouts dehydrated powder | Mini tablets | 1 tablet: 3x day; before meals | 4.2 | 1 | Not indicated |
| Daosin | Sciotec | Porcine kidney extract | Tablets | 1 tablet: 3x day; before meals | 4.2 | 1 | Not indicated |
| Naturdao | Global Mopi | Vegetable protein (pea and lentil) | Tablets | 1 tablet: before meals | 32.1 | 1 | 1,000,000 |
| Histamine Manager | Sciotec | Diamine oxidase from porcine kidney protein concentrate | Capsules with pellets | 2 capsules before the consumption of histamine-rich foods | 8.4* | 2 | 20,000 |
| Histazyme | Amy Myers | Diamine oxidase from porcine kidney protein concentrate | Capsules with pellets | 2 capsules before the consumption of histamine-rich foods | 8.4* | 2 | 20,000 |
| Histamine Blocker | Bravado Labs | Vegan (not specified) | Capsules | 1 capsule before consumption of hist-rich foods | 4 | 1 | 10,0000 |
| Histamine Digest | Bioiberica | Porcine kidney extract | Capsules | 1 tablet: before meals | 4.2 | 1 | 20,000 |
| Histamine Block Plus | Seeking Health | Porcine kidney extract | Capsules | Take 2 capsules as needed, with or without food | 1.05 | 2 | 5000 |

### d. Instrumentation

Analyses were carried out by UHPLC-MS/MS using an Agilent 1290 Infinity II UHPLC system coupled to a QqQ/MS 6470 Series, both from Agilent Technologies (Santa Clara, CA, USA). The chromatographic separation was carried out on an ACQUITY UPLC BEH HILIC (1.7 μm, 2.1 mm × 100 mm) column from Waters (Milford, MA, USA). Finally a Mikro 200 Benchtop centrifuge from Hettich (Tuttlingen, Germany); a water bath orbital shaker (Unitronic, J.P. Selecta); and an Ultra-Turrax Homogenizer (Miccra D-1, Germany) were used.

### e. Dietary supplement extraction

The supplements were ground with a mortar and pestle, followed by homogenization in 0.05 M phosphate buffer solution (pH 7.2) with an ultra-turrax and placed in a water bath orbital shaker for 1h (37 °C, 40 U/min).

### f. Sample preparation

To start the enzymatic reaction, histamine was added to each dietary supplement extract to reach a final concentration of 45 μM. Six aliquots were collected at different sampling times (0, 0.5, 1, 1.5, 2, and 3 h). Positive control samples were performed with purified DAO from Sigma (5/10 mg).

The enzymatic reactions were stopped by adding 15 μL of 2N HClO4 to the reaction mixture. An internal standard 50 μg/mL Histamine-d4 was also added.

The mixture was centrifuged at 15000 rpm for 5 minutes at 4ºC. The supernatant was diluted in a solution of 0.1 % formic acid in acetonitrile and transferred to a clean vial for LC-MS/MS analysis.

### g. Quantification of histamine

A gradient-based chromatographic separation was conducted using two phases: 100% water with 100 mM ammonium formate (pH=3) and 100% acetonitrile. The column temperature was maintained at 45 ºC and the injection volume was 2 μL.

## 3. Results and discussion

Ten over-the-counter dietary supplements, comprising seven from porcine kidneys and three from vegetable sources were analysed for their DAO activity, as presented in (Table 1).

The DAO activity of the analysed dietary supplements ranged from 0 to 0.42 mU/mg of extract. Among the tested supplements, DAOfood Plus, DAOfood, DAOfood Veg. exhibited the highest DAO activity expressed as mU/mg of extract. Upon comparison of porcine-derived DAO supplements, no statistically significant difference was observed between DAOfood and DAOfood Plus. However, the enzymatic activity of both supplements was found to be significantly higher when compared to the remaining supplements. Histamine manager, Daosin and Hystazyme also displayed some activity, but it was significantly lower (Figure 1). In contrast, the supplements Histamine digest, and Histamine block plus showed no detectable activity (Figure 1).

**Figure 1.**
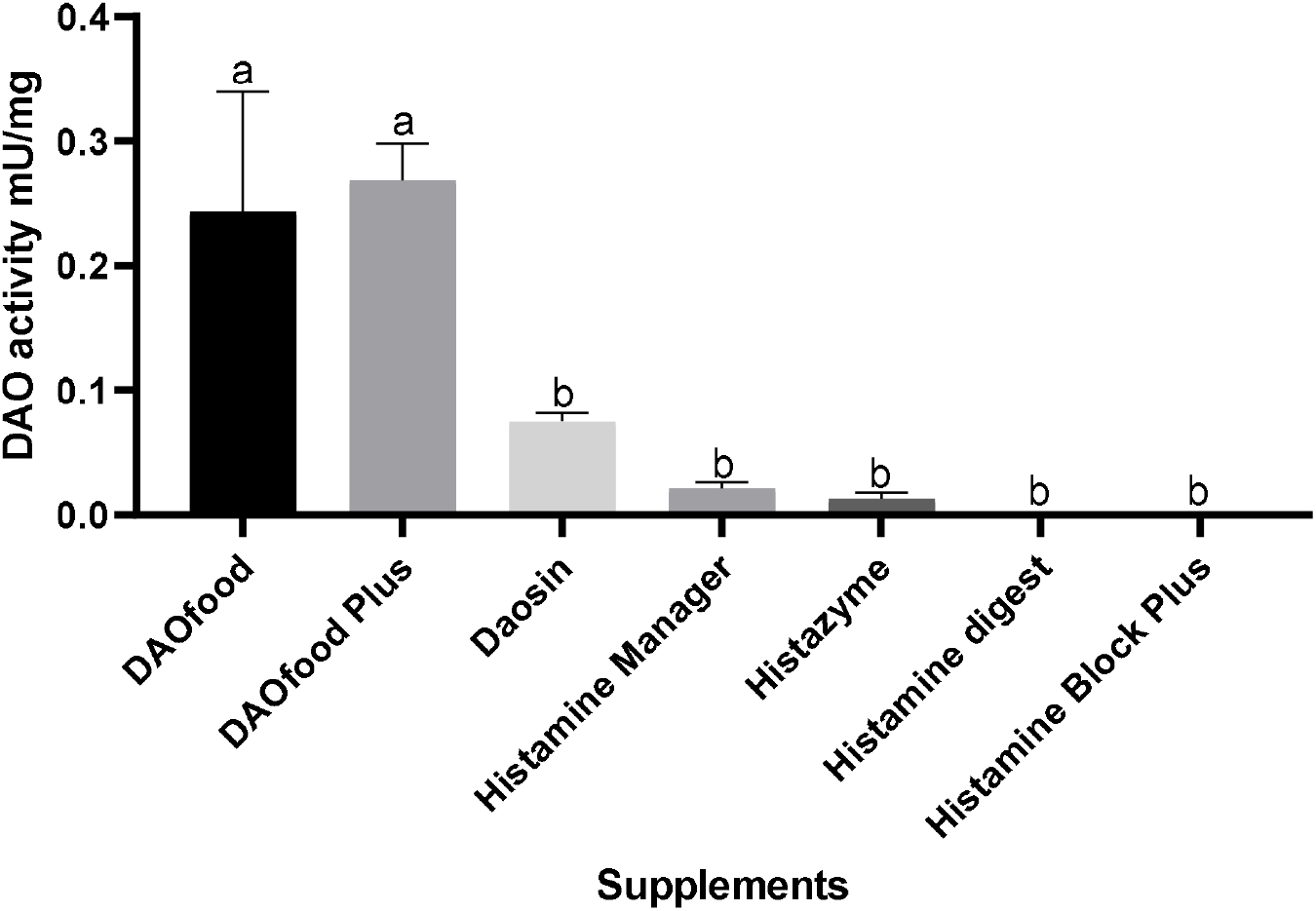
One-way ANOVA of the DAO activity detected expressed as mU/mg of extract for the DAO supplements from animal source (pvalue<0,05). *The values for the amount of protein extract rich in DAO for the products Histamine manager and Histazyme have been estimated as the label of those products (300mcg=0,3mg) refers to the content of DAO enzyme, not to the content of protein extract, like the rest of products. For ease of comparison, 0.3 mg is the estimated content of DAO in 4.2 mg (1 capsules) of protein extract as stated in EU Regulation 2017/2470(17).

When comparing plant-based DAO supplements among them, only one of the three analysed showed diamine oxidase activity (Figure 2).

**Figure 2.**
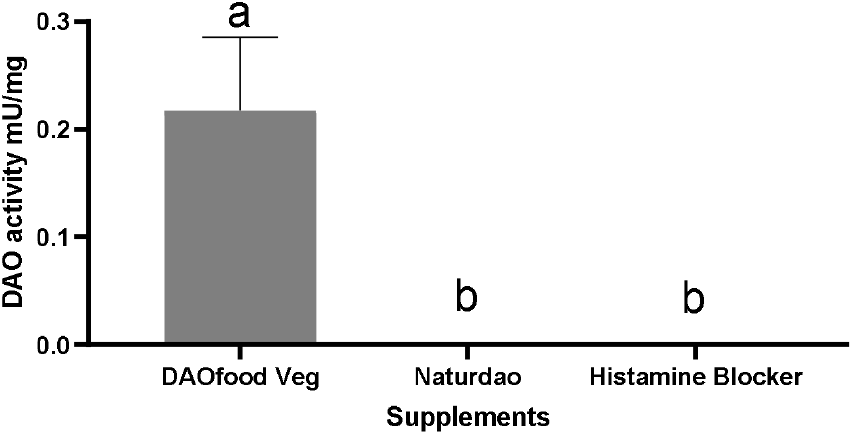
One-way ANOVA of the DAO activity detected expressed as mU/mg of extract for the DAO supplements from vegetal source (pvalue<0,01).

Within the products included in this study, eight declared the DAO activity in HDU per serving on the packaging, while two did not. Table 2 demonstrates the % of the stated activity, which ranged from 0% to 244.445% of the stated label amount. The table reveals that only two supplements contained at least 100% of the stated label claim for DAO activity, while eight had no declared activity or the activity detected did not reach the stated amount.

**Table 2.** DAO activity declared versus activity found in supplement products (n = 10). Data reported as average of two analyses. * 1 mU is equivalent to 48 000HDU of the DAO radioextraction method of analysis as determined by O.Comas-Basté_16_.

| Supplement | DAO activity label specifications (HDU/serving) | Detected DAO activity (HDU/serving)* | Detected DAO activity (mU/serving) | Relative activity % |
| --- | --- | --- | --- | --- |
| DAOfood Plus | 20,000 | 48,889 | 1.008 | 244.45 |
| DAOfood | 20,000 | 40,742 | 0.84 | 203.71 |
| DAOfood Veg | Not indicated | 30,556 | 0.63 | N/A |
| Daosin | Not indicated | 14,260 | 0.07 | N/A |
| Naturdao | 1,000,000 | 0 | 0 | 0 |
| Histamine Manager | 20,000 | 8148 | 0.168 | 40.74 |
| Histazyme | 20,000 | 4074 | 0.084 | 20.37 |
| Histamine Blocker | 10,000 | 0 | 0 | 0 |
| Histamine Digest | 20,000 | 0 | 0 | 0 |
| Histamine Block Plus | 5000 | 0 | 0 | 0 |

In addition to the findings depicted in Figure 1, an assessment of DAO activity per serving, which reflects more real-life conditions, reveals that DAOfood Plus and DAOfood exhibit activity levels that surpass the minimum required amounts, while Histazyme and Histamine manager and Daosin, exhibit lower activity levels when the amount per serving is taken into consideration.

Most DAO dietary supplements sold contained less DAO activity than the stated label amount. Several reasons could account for that phenom. One reason may be fluctuations in the DAO source, as all the supplements studied obtain their DAO activity from natural sources, which are subject to the heterogeneity of biology and natural products. Animals and plants from different sources, such as different breeds or raising locations, may have different DAO activity levels based on raising regimens(18–20). Moreover, the lack of standardization of extract production could result in high variability across different batches, degrade a part of the enzyme when produced, or deliver an enzyme with poor stability over time.

This study demonstrates the significant variability between the stated DAO activity and the one determined analytically. These findings suggest that there is still room for improvement in DAO supplements manufacturing. Individuals who are taking DAO supplementation should prioritize trustworthy brands with robust standardization protocols to ensure the efficacy of the supplement for the proper management of histamine metabolism. Moreover, it is essential to seek consultation with trained physicians to ensure proper guidance on the appropriate usage and dosage of DAO supplements. For healthcare practitioners, several recommendations for selecting proper supplements can be followed as consulting the analytical methods used for manufacturers (e.g., HPLC tends to be the gold standard), having acquired GMP certification and conducting clinical trials, among others (21).

## 4. Conclusions

As a result of this study, a wide range of products labelled as containing Diamine Oxidase were evaluated, but most failed to meet the desired quality standard for the correspondence between their actual DAO activity and the value reported on the labelling.

In light of the findings from this study, it is crucial that consumers exercise caution when purchasing dietary supplements containing Diamine Oxidase. Despite the different wide range of DAO supplements, a significant proportion of them exhibited lower enzymatic activity than their advertised levels. Moreover, some even lack basic information about their ingredients. This makes it difficult for consumers to know what they are putting into their bodies and can lead to unpredictable efficacy outcomes. Therefore, it is important to acquire DAO supplements from trusted companies that put effort into following strict quality control measures, even though it is not yet mandatory for dietary supplements. By doing so, consumers and Healthcare practitioners can ensure that they are getting a reliable and safe product that is consistent with the labelling information.

## Acknowledgements

We thank Dr. Antoni del Pino and Iris Samarra for their support in sample analysis.

